# Ultra-deep through-skull mouse brain imaging via the combination of skull optical clearing and three-photon microscopy

**DOI:** 10.1101/2021.12.20.473469

**Authors:** Mubin He, Dongyu Li, Zheng Zheng, Hequn Zhang, Tianxiang Wu, Weihang Geng, Zhengwu Hu, Zhe Feng, Shiyi Peng, Liang Zhu, Wang Xi, Dan Zhu, Jun Qian

## Abstract

Optical microscopy has enabled *in vivo* monitoring of brain structures and functions with high spatial resolution. However, the strong optical scattering in turbid brain tissue and skull impedes the observation of microvasculature and neuronal structures at a large depth. Herein, we proposed a strategy to overcome the influence induced by the high scattering effect of both skull and brain tissue via the combination of skull optical clearing (SOC) technique and thee-photon fluorescence microscopy (3PM). The Visible-NIR-II compatible Skull Optical Clearing Agents (VNSOCA) we applied reduced the skull scattering and water absorption in long wavelength by refractive index matching and H_2_O replacement to D_2_O respectively. 3PM with the excitation in the 1300-nm window reached 1.5 mm cerebrovascular imaging depth in cranial window. Combining the two advanced technologies together, we achieved so far the largest cerebrovascular imaging depth of 1.0 mm and neuronal imaging depth of >700 μm through intact mouse skull. Dual-channel through-skull imaging of both brain vessels and neurons was also successfully realized, giving an opportunity of non-invasively monitoring the deep brain structures and functions at single-cell level simultaneously.

## Introduction

Brain structural and functional imaging is essential for monitoring neuronal circuitry and pathogenesis of brain diseases including Alzheimer’s disease and Parkinson’s disease^1-6^. Optical imaging is optimal for its high spatial resolution, real-time performance and free of ionizing radiation. However, the strong scattering effect of biological tissues^7^ has impeded it from observing vasculature and neuronal structures at a large depth. Many technologies have been developed to reduce the scattering effect, including photoacoustic tomography^8-10^ with acoustic penetration, optical coherent tomography^11-14^ with optical heterodyne amplification, and so on. However, these modalities performed poorly in neuronal structure imaging due to the lack of specific mark to neuron, let alone neuronal dynamic imaging.

The mouse cortex is mainly protected by the skull, which exhibits strong scattering, causing a severe limitation for large-depth and high-contrast optical imaging. In fact, several technologies have been proposed to avoid the influence of optical scattering caused by the skull, including the open-skull glass window^15^, the thinned-skull cranial window^16, 17^ and the skull optical clearing window^18, 19^. The open-skull glass window, which is formed by removing a part of skull and replacing with a glass coverslip, tends to induce a series of inflammatory reactions and increase intracranial pressure^20^. The thinned-skull cranial window, which is literally formed by thinning the skull, requires operating repeatedly due to bone growth and is difficult to use^16^. However, the skull optical clearing window, which locally applies some chemical agents to the skull for refractive index matching, is safe and repeatable, without any craniotomy^18^. Previously developed skull optical clearing windows have enabled repeated imaging of the dendritic protrusions, microglia dynamics and blood capillaries, combined with optical microscopy^18, 19^. Nevertheless, most of them employed H_2_O as solution, which has large absorption coefficient in near-infrared region, especially beyond 1300 nm. This would inevitably cause adverse impact on optical imaging with near-infrared excitation or emission.

In addition, mouse brain tissue also exhibits strong scattering effect and impedes deep imaging, unable to be resolved by *in vivo* tissue optical clearing technology at present. Fortunately, three-photon fluorescence microscopy (3PM) is a typical way for deep imaging with high contrast in turbid tissue^21^. The character of long-wavelength excitation light with low tissue scattering guarantees 3PM large penetration depth, thus maintaining well focus efficiency in turbid tissue^22^. Meanwhile, high-order nonlinear optical effect of three-photon fluorescence (3PF) ensures the localized excitation, and therefore improves imaging resolution and contrast. The two acknowledged long-wavelength excitation windows for 3PM are the 1300-nm and 1700-nm spectral ranges^23^, both of which have long attenuation length at brain tissue. Among the two excitation windows, the 1300-nm window is favored for its lower tissue absorption induced heating effect. Moreover, considering universal commercial fluorescent probes for neuronal labeling, the 1300-nm window is more commonly used in 3PF neuronal structural and functional imaging^24, 25^, and more capable of simultaneous multi-color imaging^26^. However, 3PM using the 1300-nm excitation under the intact skull currently achieved only ∼300 μm neuronal imaging depth^25^ and 510 μm cerebrovascular imaging depth^25^ owing to the low excitation efficiency.

Herein, we combined *in vivo* skull optical clearing (SOC) technique and 1300-nm excited 3PM to overcome the scattering and absorption of the skull as well as brain during deep-cortical visualizing. Skull scattering reduction could improve the performance of 3PM effectively according to Monte Carlo simulation results. The Visible-NIR-II compatible Skull Optical Clearing Agents (VNSOCA) we applied introduced D_2_O solution to replace H_2_O solution. Therefore, the light scattering and absorption of skull in the 1300-nm window were both reduced effectively. Afterwards, we realized largest 3PF through-skull cerebrovascular imaging depth (1.0 mm), assisted by a kind of fluorescent probe with large three-photon absorption cross section. Meanwhile, 3PF through-skull neuronal imaging also improved to >700 μm depth. With prominently reduced scattering effect and near-infrared absorption of skull, 3PM in the 1300-nm window achieved dual-channel deep imaging of both brain vessels and neurons under the intact skull for the first time.

## Results

### Skull scattering reduction improved performance of through-skull 3PM

Skull scattering is one of the major barriers that constrains the through-skull imaging quality. And SOC technique could reduce the skull scattering effectively. Herein, we performed Monte Carlo simulation to investigate how SOC would enhance the performance of three-photon excitation due to the better focusing inside brain tissue. Fig. 1a shows the simulated distribution of photons when the 1300-nm light was focused on the brain tissue through skull with the scattering of skull decreasing. Fig. 1b states the situation when a 1700-nm light was focused on the brain tissue. It was obvious that reducing the skull scattering would effectively reduce the out-of-focus light both for 1300-nm window and 1700-nm window. In addition, quantitative analysis demonstrated that, both the photon intensity and photon density at the focus point were getting better as the skull scattering coefficient decreased in the 1300-nm window (Fig. 1c and 1d) and the 1700-nm window (Fig. 1e and 1f). In addition, we simulated the amount of signal intensity left at the skull surface after penetrating through brain tissue and skull when the emission was generated at certain depth of brain. As shown in in Supplementary Figure 1, the signal photons showed similar distribution on the surface as the skull scattering coefficient decreased. However, the overall photons penetrating successfully to the surface obviously increased as the skull scattering coefficient decreased (Supplementary Figure 2). Simulation results above verified the skull scattering reduction induced by SOC could promisingly increase the photon density of the three-photon excitation light at focus point, and decrease the attenuation of the generated three-photon emission signals before detected.

**Fig. 1.**
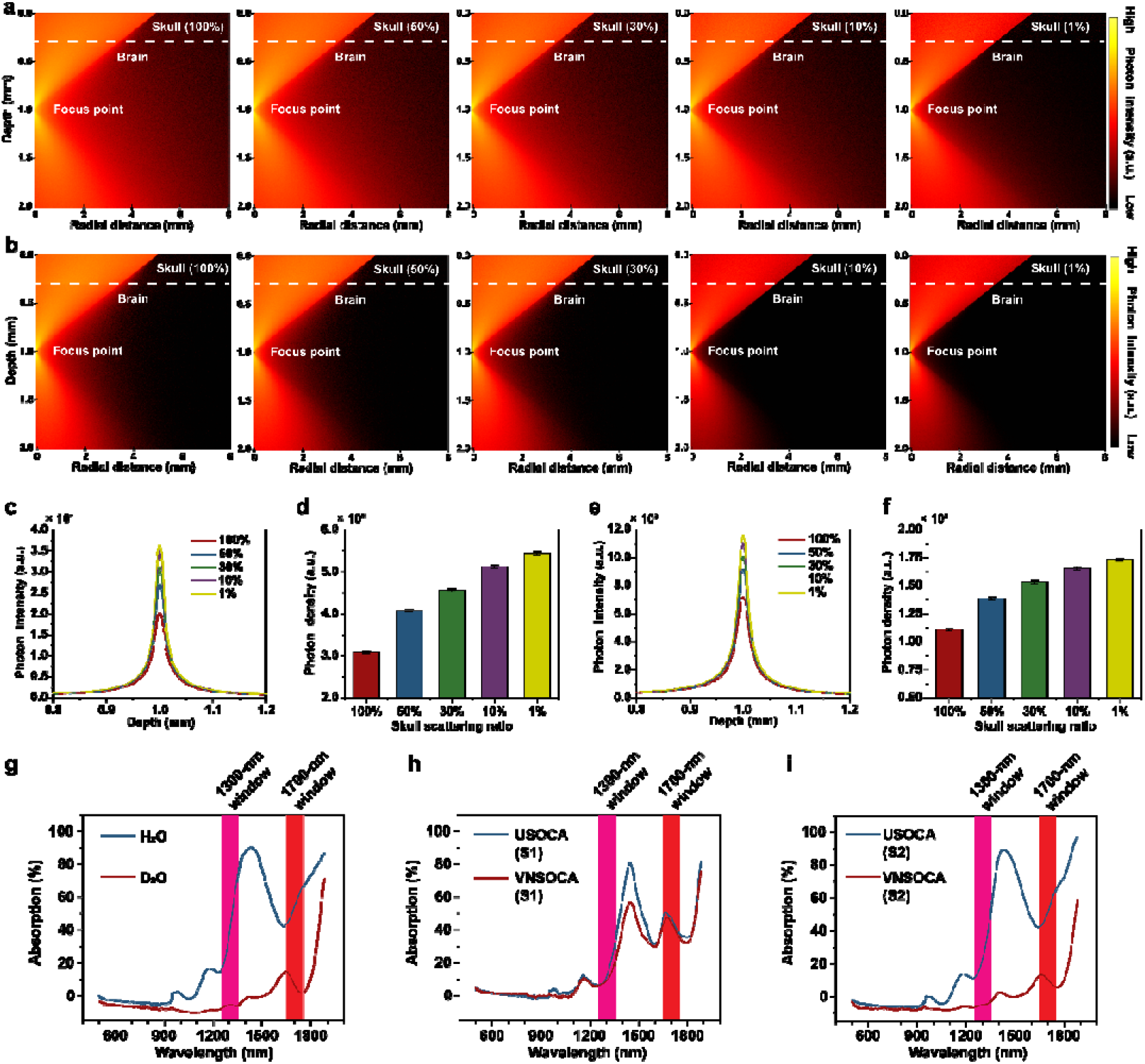
Skull scattering reduction improved performance of through-skull 3PM. **a, b** R-Z plane photon distribution simulation of excitation light in the 1300-nm window (a) and the 1700-nm window (b) penetrating through skull and brain tissue with different remaining skull scattering ratio (100% to 1%). Focal depth = 1 mm. **c, e** Photon intensity (number of photons at focal plane) profiles along the depth (Radial distance = 0 mm) from 0.8 mm to 1.2 mm with different remaining skull scattering ratios (100% to 1%) in the 1300-nm window (c) and the 1700-nm window (e). **d, f** Photon density (number of photons divided by focal spot area at focal plane) at the focal point with different remaining skull scattering ratio (100% to 1%) in the 1300-nm window (d) and the 1700-nm window (f). **g** Absorption spectra of H_2_O and deuterium oxide (D_2_O). **h, i** Absorption spectra of solution 1 (S1) and immersed solution 2 (S2) of urea-based skull optical clearing agents (USOCA) and visible-NIR-II compatible skull optical clearing agents (VNSOCA). (Optical length = 1 mm).

Light absorption by water in near-infrared region was another factor that should be considered while applying the skull optical clearing agent (SOCA) to 3PM. Since deuterium oxide (D_2_O) has been applied as an immersion medium in near-infrared imaging for its reduced absorption in near-infrared region^27^ (Fig. 1g), in this work we prepared visible-NIR-II compatible skull optical clearing agents (VNSOCA) which replaced H_2_O in urea-based skull optical clearing agents (USOCA) with deuteroxide solution. Then we used Monte Carlo simulation to compare two SOCAs to investigate the impact of absorption of the SOCA (Fig. 1h and 1i) on 3PM. As shown in Supplementary Figure 3, after SOC treatment, solution 2 (S2) of the SOC was remained on the skull for imaging. Therefore, simulations were performed to investigate the 1300-nm and 1700-nm light focusing ability when S2 of USOCA and VNSOCA covered skull. As shown in Supplementary Figure 4, compared with USOCA, VNOSCA can increase the 1300-nm photon density at the focal point by 1.25 times. Similarly, as shown in Supplementary Figure 5, VNSOCA can increase the 1700-nm photon density at the focal point by 2 times. However, in the case of VNSOCA, the absolute value of 1700-nm photon density was lower than that of 1300-nm photon density at the focal point (1.4_×_10^4^ VS 3.1_×_10^4^), due to stronger brain tissue absorption.

Furthermore, we also performed *ex vivo* 3PM test to verify the skull clearing effect by VNSOCA. A kind of lab-synthesized aggregation-induced emission (AIE) nanoparticle (NP), DCBT NP (the details will be discussed later), and a lab-built three-photon microscopic system (supplementary Figure 6), were used for the test. The results indicated that, covered by an intact skull, the 3PF signals of DCBT NPs-filled capillary were rather weak. On the contrary, when the skull was treated by VNSOCA, the signals were remarkably increased (Supplementary Figure 7). In addition, VNSOCA treatment also increased the signal-to-background ratio (SBR, supplementary Figure 8) and the resolving power (supplementary Figure 9) of 3PM.

Overall, VNOSCA assisted skull optical clearing could reduce both light absorption by water in near-infrared region and skull scattering, therefore enhanced the performance of through-skull 3PM.

### The advantages of 1300-nm excitation window over 1700-nm excitation window

The advantages of long excitation wavelength and high-order nonlinear optical confinement attributed to 3PM were effective ways to overcome the scattering effect in deep brain imaging. The greatest advantage of the 1300-nm excitation window against the 1700-nm excitation window is its much lower tissue absorption. The Monte Carlo simulation of temperature rising caused by tissue absorption of excitation light was shown in Fig. 2a-b and supplementary Figure 10. Compared with 1700-nm window, temperature rising in the 1300-nm window was obviously milder. Also, the temperature distribution in the 1300-nm window was more dispersive, avoiding local overheating at focused point. On the contrary, 1700-nm light would cause a sharp temperature rise near the focus, where overheated damage tended to happen easier.

**Fig. 2.**
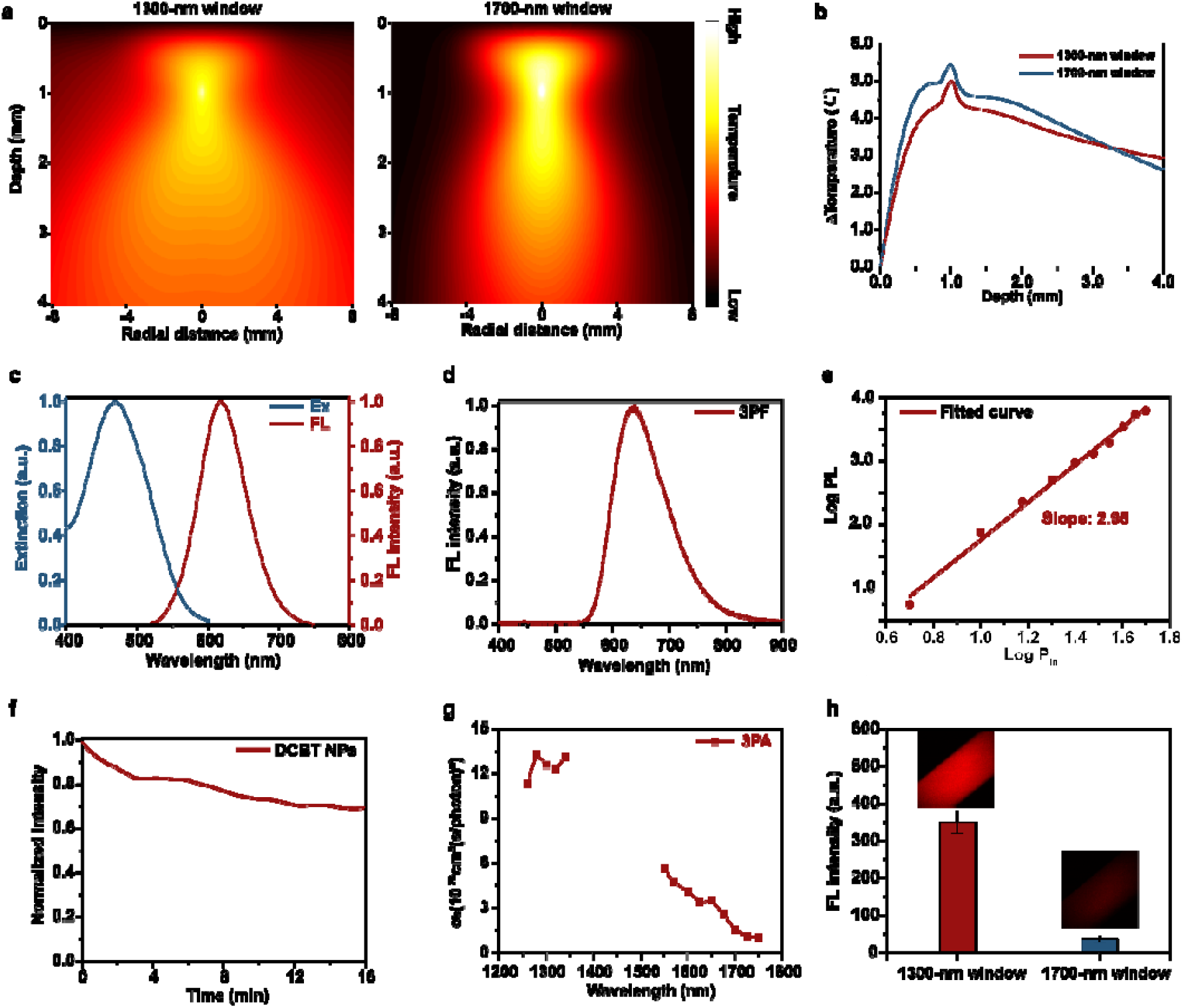
The advantages of the 1300-nm excitation window over 1700-nm excitation window. **a** Monte Carlo simulation of temperature rising distribution caused by tissue absorption in the 1300-nm window and 1700-nm window. (Focal depth = 1 mm). **b** Temperature rising distribution line versus depth at the position of 0 mm radial distance. **c** Normalized extinction and PL spectra of DCBT NPs. **d** 3PF spectrum of DCBT NPs under the excitation of 1300 nm fs laser. **e** Power dependence of fluorescence intensity of DCBT NPs under the 1300 nm fs excitation intensity. **f** Photostability of DCBT NPs under the continuous irradiation of 1300 nm fs laser (Average power: 7 mW). **g** Measured three-photon absorption cross-sections of DCBT NPs from 1260 nm to 1750 nm. **h** 3PF intensity of DCBT NPs under the excitation of 1300-nm window and 1700-nm window. Insert: 3PF images of DCBT NPs filled in the glass capillary.

Another advantage of the 1300-nm excitation window was that compared with the 1700-nm excitation window, the 1300-nm excitation window should obtain higher excitation efficiency and fluorescence intensity for its shorter wavelength according to the multi-photon fluorescence photon flux formula^28^. However, the low three-photon absorption cross-section (σ3) of most fluorescence probes restricted their 3PF intensity from further strengthening. Fortunately, we currently developed an AIE probe named DCBT with large σ3 which was realized by the intra- and intermolecular synergistic engineering. The AIE character of DCBT endows it with bright fluorescence and high photostability in the aggregated nanoparticles. Considering those unique properties, DCBT was introduced in this study and its σ3 in the 1300-nm and 1700-nm excitation windows were evaluated. As shown in supplementary Figure 11, DCBT were encapsulated by F-127, which was approved by the US Food and Drug Administration (FDA), to form amphipathic organic nanoparticles. The results of dynamic light scattering (DLS, 36.2 ± 1.9 nm, Zetasizer Nano-ZS) and scanning transmission electron microscopy (STEM, 32.3 ± 3.88 nm, tecnai Spirit) in supplementary Figure 12 showed the micellar systems were uniformly distributed. The extinction and photoluminescence (PL) spectra were displayed in Fig. 2c. The quantum yield of DCBT NPs was measured as 3.17%. Afterwards, we measured its nonlinear emission spectrum. As shown in Fig. 2d, under the 1300-nm fs excitation, DCBT NPs generated red emission with a peak at ∼640 nm. In addition, the dependence relationship between the nonlinear fluorescence intensity and the excitation light intensity was measured. As shown in Fig. 2e, the logarithm of the emission intensity against that of the excitation intensity showed a linear slope of approximately 2.95, indicating a major three-photon absorption process under 1300-nm fs excitation. Moreover, 3PF intensity decreased by only 30% over 16 minutes of continuous irradiation (Fig. 2f), suggesting the promising photostability of DCBT NPs for 3PM.

Furthermore, σ_3_ of DCBT in aggregation at various wavelengths were measured, since it was the major feature manifesting the 3PM capability of an optical probe. Results showed that, the σ_3_ of DCBT at 1300 nm was much larger than that at 1700 nm (Fig. 2g). The largest σ_3_ at 1300 nm within the test range was calculated to be 1.26 × 10^−77^ cm^6^ s^2^, which is the highest value among the reported organic probes^26, 29^.

Hence, 3PF intensity excited at the 1300-nm wavelength was ∼10 times higher than that excited at the 1700-nm wavelength (Fig. 2h). Therefore, with the help of the improved σ3 of fluorescence probe, *in vivo* 3PF imaging in the 1300-nm window would be more suitable for the observation of brain tissue with the high scattering.

### *In vivo* deep 3PF brain vasculature imaging of the mouse with the open-skull glass window

With the assistance of DCBT NPs and utilization of open-skull glass window (Fig. 3a), we adopted 3PM to perform ultra-deep visualization of mouse cerebral vessels. The mouse was imaged after injected with DCBT NPs dispersion (1.5 mg/mL, 0.2 mL). Fig. 3b showed 3D reconstruction of cerebral vessels under the excitation of 1700-nm window and 1300-nm window. The imaging depth of 3PM using the 1700-nm window only reached 1.1 mm (Supplementary Figure 13) due to the low three-photon action cross-section. In contrast, 3PM performed with the 1300-nm window could reach 1.5-mm imaging depth. This imaging depth has already passed through the white matter layer and reached the hippocampus regions, and broke the previously reported organic probe-assisted 3PF imaging depth record (1.4 mm^29^) using the 1300-nm window.

**Fig. 3.**
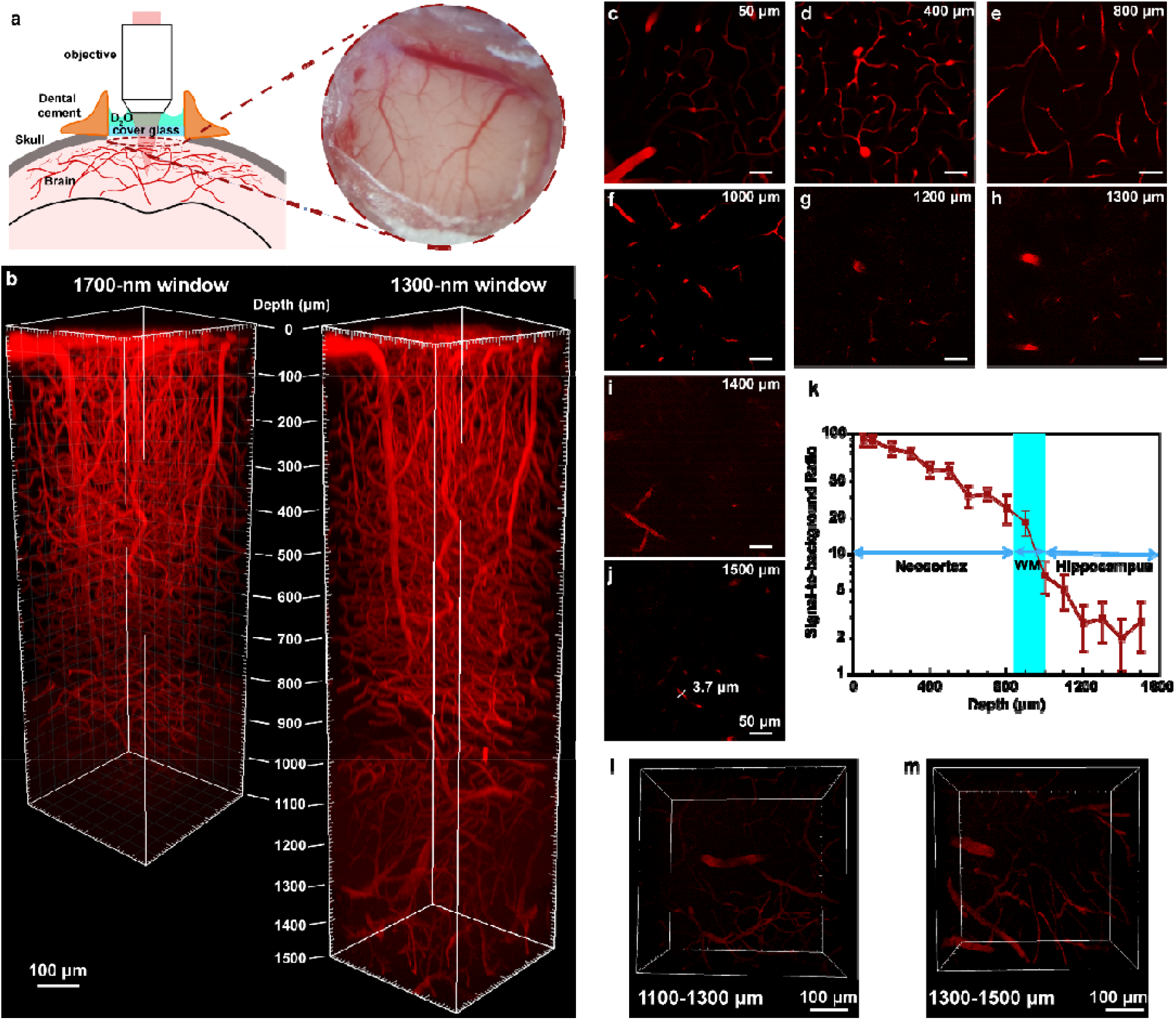
*In vivo* deep 3PF brain vasculature imaging of the mouse with the open-skull glass window. **a** Schematic of mouse head imaging with the open-skull window. The bright field picture of mouse brain was shown at right position. **b** 3D reconstruction of the cerebrovascular imaging under 3PM excited using the 1700-nm window (0-1100 μm) and 1300-nm window (0-1500 μm). **c-j** 3PF imaging of the mouse brain vasculature at various depths excited in the 1300-nm window **k** The Signal-to-background ratio (SBR) as a function of imaging depth. The white matter (WM) region of the mouse brain, where the SBR decreased sharply, is indicated in blue. **l-m** 3D reconstruction of the cerebrovascular imaging in 1100-1300 μm and 1300-1500 μm depth.

Brain vessels at various depth was shown in Fig. 3c-j. The tiny blood capillary with a size of 3.7 μm could be distinguished at the 1.5 mm depth (Fig. 3j and Supplementary Figure 14). Images at neocortex layer maintained extremely high SBR due to the background-free feature of 3PM as shown in Fig. 3k. Some large blood vessels were observed from white matter layer (800 μm in Fig. 3e) to hippocampus layer (1000 μm in Fig. 3f), which was consistent with the results in previous 3PF deep imaging works^30-32^. Because of the higher scattering effect of white matter, SBR appeared to decrease when the region of interest (ROI) travelled through it (Fig. 3k). Such situation is similar to what was observed in long-wavelength excited confocal fluorescence imaging^33^.

Nevertheless, even with relatively low SBR (∼2), the vessels were still distinguishable. 3D reconstruction of brain vessels in the 1.1-1.3 and 1.3-1.5 mm was displayed in Fig. 3l-m. Large blood vessels extended up to 1.5 mm below was displayed clearly. However, it was the working distance of the objective (2 mm) that constrained imaging depth available in our experiment. As demonstrated by the results above, the scattering effect of brain was effectively overcome by 3PM with the improved σ3 of DCBT NPs in the 1300-nm excitation window.

### *In vivo* deep brain 3PM under the 1300-nm excitation window with the optical clearing window

Combining the SOC technique and 3PM together, we performed 3PF microscopic imaging of the mouse with the skull optical clearing window. The schematic of mouse head imaging was displayed in Fig. 4a. The typical bright field view of the mouse brain vasculature after skull clearing in Fig. 4b showed more apparent than that before skull clearing. Intensity analysis of vasculature under the VNSOCA treated skull in Fig. 4c demonstrated four distinct brain vessels, which were almost invisible under the untreated turbid skull. Afterwards, the 3PF cerebrovascular imaging before and after skull clearing were carried out on mouse injected with DCBT NPs. As Fig. 4d-g showed, 3PF imaging performed through the original skull had only reached 550 μm, which was comparable with the previously reported imaging depth^25^. On the contrary, after VNSOCA treatment on the skull to decrease its scattering, 3PF imaging of the same field of view showed enhanced signal intensity and optimized SBR (Fig. 4h-o), especially at large depth. The final 3PF imaging depth reached 1.0 mm through the skull optical clearing window (Fig. 4p-s). The small blood vessel with a diameter of 4.5 μm at 700-μm depth was clearly visualized (Fig. 4p, t). Moreover, a 10.4-μm vessel at 1000 μm depth could also be distinguished (Fig. 4s, u). 3D reconstruction of brain vessels from 0 to 1000 μm was displayed in Fig. 4v. To the best of our knowledge, our 3PM achieved the largest millimeter-level imaging depth under the intact skull, thanks to the reduced skull scattering induced by VNSOCA and reduced brain tissue scattering overcome by 3PF.

**Fig. 4.**
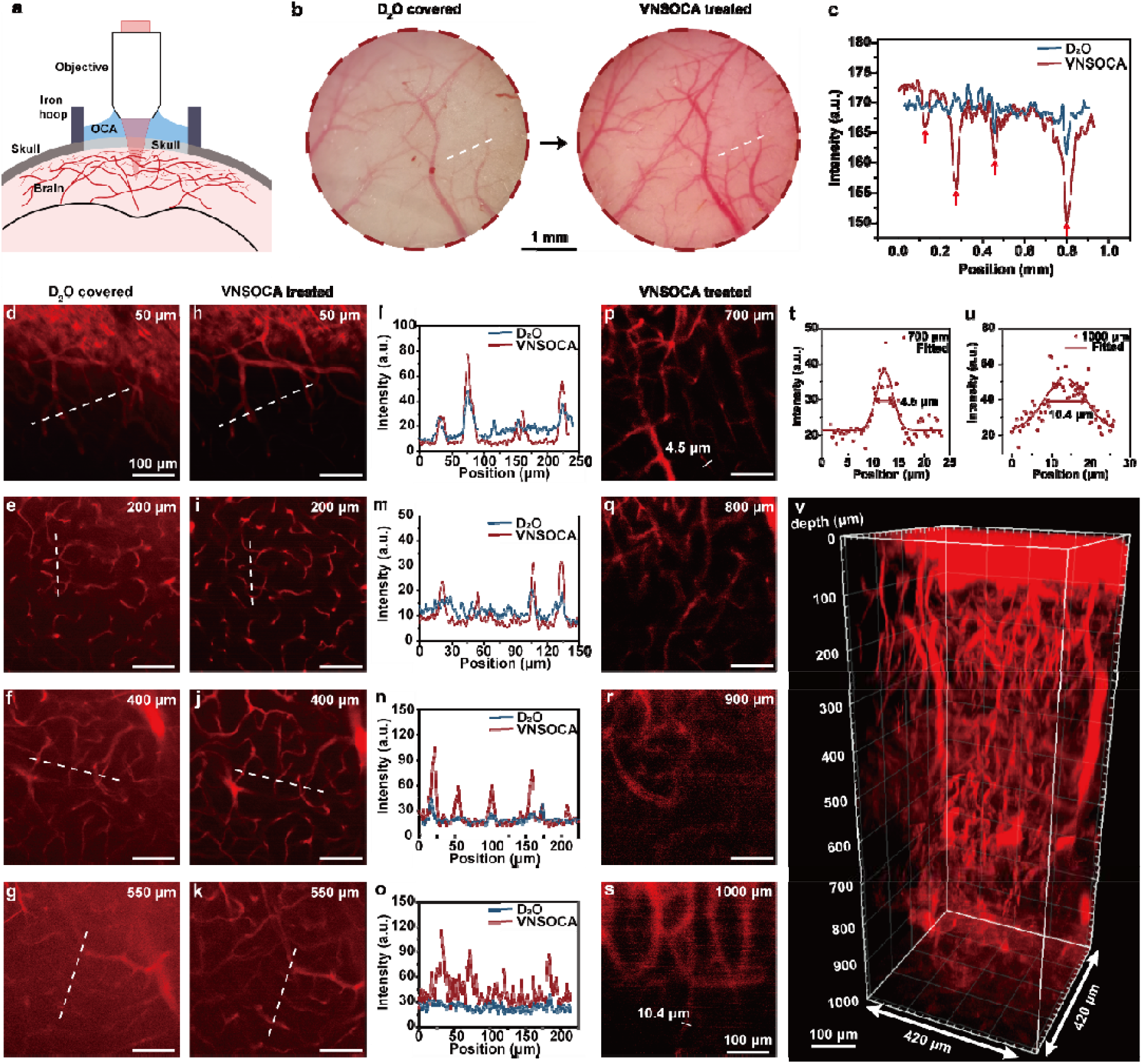
*In vivo* deep 3PF brain vasculature imaging of the mouse with the optical clearing window. **a** Schematic of mouse head imaging with the optical clearing window. **b** Typical bright field pictures of the mouse brain vasculature before and after skull clearing. **c** Intensity profiles along the white dashed lines across the vasculature in (b). The arrows indicated the positions of vessels. **d-g** 3PF imaging of the mouse brain vasculature at various depths with original skull. **h-k** 3PF imaging of the mouse brain vasculature at various depths after skull clearing. **l-o** Intensity profiles along the white dashed lines across the brain vasculature in (d)-(k) respectively. **p-s** Deep 3PF images of the mouse brain vasculature after skull clearing. **t-u** FWHM analysis of the brain vasculature at the depths of 700 μm (t) and 1000 μm (u). **v** 3D reconstruction of the cerebrovascular imaging with the optical clearing window (0-1000 μm).

Apart from cerebrovascular imaging, visualization of mouse brain neurons at neocortex layers was also conducted under the intact skull. As shown in Fig. 5a-d, not only the signal intensity, but also the SBR and resolution of neuronal dendrite were considerably enhanced at shallow depth after skull optical clearing. Fig. 5e-f showed that the skull optical clearing treatment introduced a 6-time improvement of the 3PF intensity, and compressed the half-height full width of an imaged dendrite from 3.1 microns to 1.2 microns at 25-μm depth. More abundant dendrites emerged at the depths of 75 μm and 160 μm after skull optical clearing compared with those observed through original skull (Fig. 5g-j).

**Fig. 5.**
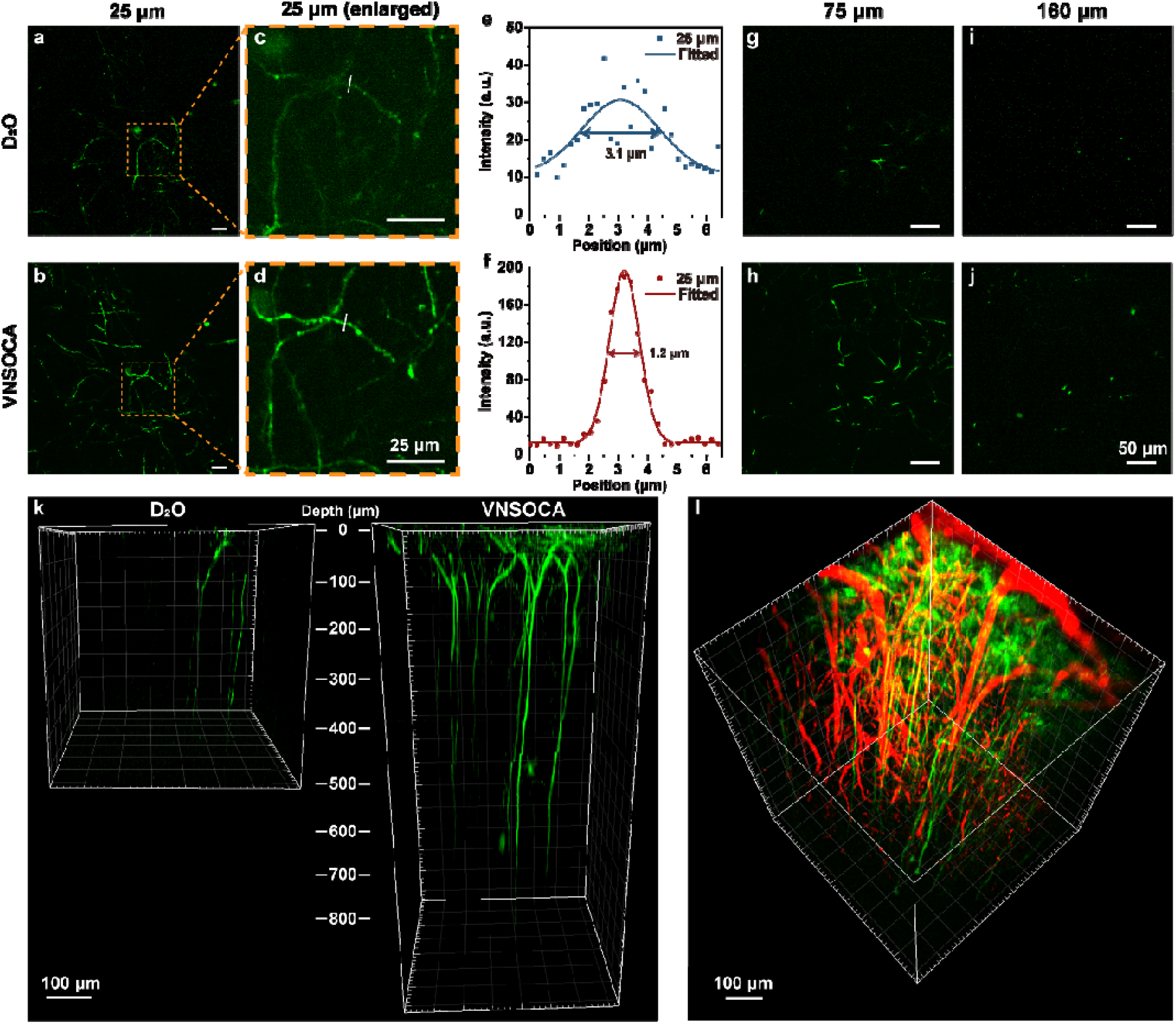
*In vivo* deep 3PF brain neuron imaging and neuron-vessels dual channel imaging of the mouse with the optical clearing window. **a-b** 3PF images of the mouse brain neurons at 25-μm depth before and after skull clearing. **c-d** Enlarged 3PF images of the mouse brain neurons in (a) and (b). **e-f** Intensity and FWHM analysis of the brain neurons in (c) and (d). **g-j** 3PF images of the mouse brain neurons before and after skull clearing at depth of 75 μm and 160 μm. **k** 3D reconstruction of the 3PF neuronal imaging through the original skull (0-400 μm) and the skull optical clearing window (0-800 μm). **l** 3D reconstruction of the neuron-vessels dual channel imaging of the mouse with the optical clearing window (0-600 μm).

Thanks to the strengthened signal level and improved image contrast after skull clearing, we were able to track the deep axons of *Thy-1-GFP* neurons from 250 to 700 μm (Fig. 5k and supplementary Figure 15). The vivid *Thy-1-GFP* neuron structures were visualized clearly after skull optical clearing, with improved imaging depth up to >700 μm. In virtue of the red emission of DCBT NPs under 1300-nm three-photon excitation, multicolor fluorescence imaging was also carried out. The 3D reconstruction in Fig. 5l clearly displayed the merged image of the *Thy-1-GFP* neurons and DCBT NPs-labeled vessels from 0 to 600 μm. So far as we know, this is the first reported 3PF deep and simultaneous imaging of brain vessels and neurons under the intact skull.

## Discussion

We proposed a solution to the high scattering effect of both skull and brain tissue through the combination of SOC technique and 3PM, and therefore realized ultra-deep cerebrovascular and neuronal imaging under intact skull. While previous skull optical clearing agents were mainly applied in two-photon fluorescence microscopy, overlooking large light absorption by water at long excitation wavelength, the VNSOCA we prepared could efficiently reduce both scattering and absorption coefficients of skull during 3PM process (Fig. 1). We also proposed a kind of AIE NPs with large three-photon absorption cross-sections in the 1300-nm excitation window (Fig. 2). Meanwhile, we confirmed its great 3PF performance on the mouse brain imaging with an open-skull glass window, achieving 1.5-mm imaging depth that was the largest one ever known, by employing the 1300-nm excitation window (Fig. 3). Such imaging depth reached hippocampal region, and was almost comparable to the depth achieved by the 3PM excited in the 1700-nm window^31^. In addition, 3PM on the mouse’s brain with intact skull combined with SOC technique enabled 1-mm cerebrovascular imaging depth (Fig. 4), which broke the through-skull 3PM depth record using the 1700-nm excitation window^34^. Finally, the 3PF through-skull imaging depth of neuron achieved over 700 μm. In addition, dual-channel deep imaging of both vessels and neurons under intact skull was also realized thanks to the reduced scattering of skull and brain tissue (Fig. 5).

The combination of the two techniques is quite reasonable. On the one hand, despite that 3PM alone could reduce the skull scattering effect to a certain extent, the efficiency is not enough owing to the extremely higher scattering of skull compared with the brain tissue^7^. Our previously reported 3PM on the mouse brain with intact skull reached only 400-μm^35, 36^ depth in cerebrovascular structure imaging with excitation wavelength at 1550 nm. On the other hand, although *in vivo* skull SOC could overcome the skull scattering, itself was unable to conquer the scattering effect of brain tissue. For instance, 2PM through the optical skull clearing window only reached ∼300-μm depth in the cortex^18^. Therefore, the combination of SOC technique and 3PM is a rather necessary solution to through-skull ultra-deep brain imaging. Accordingly, we realized 1000-μm 3PF cerebrovascular imaging and over 700 μm 3PF neuronal imaging of mouse brain with intact skull.

Even with the assistance of SOC technique and 1300-nm excited 3PM, ideal optical probes are still important for the final imaging performance. Although 3PM excited in the 1300-nm window has lower heating effect and higher excitation efficiency compared with that in the 1700-nm window, the stronger scattering in the 1300-nm window still constrained its imaging depth ever reported. As a solution, we proposed a new kind of AIE NPs with extraordinarily large σ3 at 1300 nm to improve the brightness of 3PF, and eventually achieved 1.5-mm 3PM depth with open-skull glass window, which was the deepest one in the 1300-nm window.

The scattering reduction strategy presented here opens the opportunity for *in vivo* ultra-deep brain 3PM under the intact skull, where the imaging depth was nearly comparable with 3PM through the cranial window^37^. In addition, the 1300-nm excitation realized multi-color 3PM, making it possible for *in vivo* observation of molecular and cellular interactions in the brain under intact skull over a large depth in future. Moreover, the SOC procedure presented here is safe and easy-handling, not as difficult as craniotomy. Therefore, 3PM based on the skull optical clearing window can be widely used in various situations. We believe that *in vivo* 3PF imaging of hippocampus region under the intact skull of mouse brain will be realized with further reduced scattering effect of both brain tissue and skull, as well as improved 3PF probes. The combination of SOC technique and 3PM will contribute to faster and deeper imaging in scattering tissues including skull and brain.

## Methods

### Monte Carlo simulation of NIR photon penetration in skull and brain tissue

The Monte Carlo method was utilized to simulate the propagation of light beams in biological tissues. There are two layers of tissue in the simulation, representing the mouse skull and mouse brain tissue, respectively. We rasterized the tissue to record the scattering and absorption events of photons. In the simulation of focusing light in the tissue, photons enter the immersion media from the air, pass through the skull and focus on a specific depth in brain layer. The position of photon is randomly set within the beam radius. Meanwhile, the focal point was set at 1 mm depth in the skull and brain tissue after refracting in multiple tissue layers. Absorption matrix is formed to record absorbed energy of photons in tissue. The temperature profile of brain can be calculated by solving the thermal diffusion equation with absorption distribution. We set the refractive index of the skull and brain tissue as 1.369, and the scattering anisotropy factor as 0.98. The absorption coefficient of skull and brain tissue was set according to the water absorption coefficient in 1300 nm and 1700 nm. The reduced scattering coefficient of brain tissue was calculated using the following formula^7^: 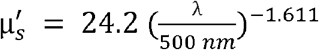. Thus, the brain tissue scattering coefficient in Fig. 1 was set as 5.192 cm^-1^ in the 1300-nm window and 3.370 cm^-1^ in the 1700-nm window respectively. The reduced scattering coefficient of skull was calculated using the following formula^7^: 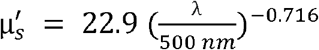. Accordingly, the remaining skull scattering coefficient in Fig. 1 was set as 11.554 cm^-1^ (100%), 5.777 cm^-1^ (50%), 3.466 cm^-1^ (30%), 1.155 cm^-1^ (10%), 0.116 cm^-1^ (1%) in the 1300-nm window and 9.534 cm^-1^ (100%), 4.767 cm^-1^ (50%), 2.860 cm^-1^ (30%), 0.953 cm^-1^ (10%), 0.095 cm^-1^ (1%) in the 1700-nm window respectively.

### Preparation and vis-NIR-II absorption measurement of optical clearing agents

The visible-NIR-II compatible skull optical clearing agents (VNSOCA) contain a saturated supernatant solution of urea and ethanol (named S1), and a high-concentration sodium dodecyl benzenesulfonate (named S2). To prepare S1, 75% (vol/vol) ethanol in deuteroxide solution was dropped onto the excessive urea and mixed together in a beaker. The mixture was stirred for 10 min and stayed for 15 min to dissolve urea fully. The synthesized S1 was then obtained after the supernatant solution was transferred. As to S2, the sodium dodecyl benzenesulfonate (mass = 5 g) and 0.7 M NaOH in deuteroxide solution (volume = 24 mL) were mixed together, under the condition of 7.2-8 pH value. Finally, both S1 and S2 solution of VNSOCA were sealed and stored at room temperature. The urea-based skull optical clearing agents (USOCA) were prepared under the similar procedure, apart from deuteroxide replaced by deionized water. The absorption spectra of both VNSOCA and USCOA were measured from 500-1880 nm with two spectrophotometers (PG2000, Ideaoptics Instruments and NIR2200-Px, Ideaoptics Instruments).

### Three-photon fluorescence microscopic system

Three-photon fluorescence microscopic system included two major parts, a non-collinear optical parametric amplifier (NOPA) with wavelength-tunable femtosecond (fs) laser output and a commercial Bruker scanning microscope. The NOPA system included a 1030 nm fs pump laser (Spectra-Physics, Spirit) and an OPA system (Spectra-Physics, NOPA-VISIR). 1300-nm fs laser beam (115 fs, 1 MHz) and 1700-nm fs laser beam (203 fs, 1 MHz) were obtained through two amplification stages in NOPA-VISIR and introduced into the scanning microscope as excitation source. The excitation beam was focused on the sample through the objective (XLPLN25XWMP2, Olympus, NA=1.05), and the excited three-photon fluorescent signals were then collected by a GaAs PMT (H7422-40, Hamamatsu), after reflected by a 700 nm DMLP and passing through a 560 nm DMLP. The filters for red channel and green channel were FF02-641/75 (Semrock) and FF02-525/40 (Semrock).

### Optical Characterization of DCBT NPs

The absorption spectra and fluorescence spectra of DCBT NPs were measured on a UV−vis scanning spectrophotometer (UV-2550, Shimadzu, Japan) and an optical fiber spectrometer (PG2000, Ideaoptics Instruments). The photoluminescence of DCBP NPs under 1300 nm fs excitation was carried out as following. The fs laser beam traveled through a focal lens (f= 30 mm) and was focused on a cuvette, which contained the dispersion of DCBT NPs. The three-photon fluorescence signal was recorded with an optical fiber spectrometer (PG2000, Ideaoptics Instruments) through an objective (XLPLN25XWMP2, Olympus, NA=1.05). The fluorescence quantum yield of DCBT NPs was measured via the comparative method with Rhodamine 6G in DI water as reference. Details were described in our previous work^38^.

### P-I relationship measurement and three-photon absorption cross section measurement

The DCBT NPs were contained in the glass capillary and imaged under the three-photon fluorescence microscopic system. The 3PF images under various excitation powers were recorded. The intensity of three-photon fluorescence (I) vs the average power of 1300 nm fs laser (P) was plotted to determine the P-I relationship of DCBT NPs.

To obtain the three-photon absorption cross section of DCBT NPs, the comparison method was applied. The three-photon absorption cross section of TPATCN NPs at 1550 nm^38^ was selected as the reference. DCBT NPs and TPATCN NPs in aqueous medium were excited by the 1550 nm femtosecond laser, and their three-photon fluorescence signals were collected by a photomultiplier tube (PMT). The mean three-photon fluorescence intensities were calculated by ImageJ. The σ3 value of DCBT NPs was calculated by the following equation:

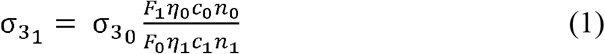

Where *F* is the three-photon fluorescence intensity, *η* is the fluorescence quantum yield, *c* is the molar concentration of sample, *n* is the refractive index of the solvent, and the subscripts 1 and 0 represent DCBT NPs and TPATCN NPs, respectively.

Three-photon absorption cross section of DCBT NPs at other excitation wavelengths were referenced to the value at 1550 nm measured above and calculated by the following equation:

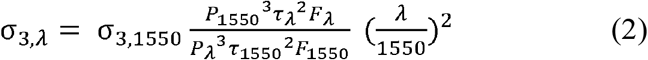

Where *σ*_*3λ*_ is the wavelength dependent 3-photon absorption cross section, *σ*_*3,1550*_ is *σ*_*3*_ of DCBT NPs at 1550 nm, *P*_*1550*_ and *P*_*λ*_ are the measured excitation powers on the sample, *τ*_*1550*_ and *τ*_*λ*_ are the measured pulse widths on the sample, and *F*_*1550*_ and *F*_*λ*_ are the measured three-photon fluorescence intensities with excitation at 1550 nm and other wavelengths, respectively.

### Ethical Approval

All animal experiments performed in this study were conducted strictly in compliance with the ethical standards of the Institutional Ethical Committee of Animal Experimentation of Zhejiang University.

### *In vivo* three-photon fluorescence microscopic cerebrovascular imaging of the mouse with the cranial window

The C57 mouse (male, 8-10 weeks old) was anesthetized by pentobarbital sodium (0.14 mL, mass concentration = 1%), and a cranial window with diameter of around 4 mm was produced by removing the scalp and a small piece of skull. After injected with DCBT NPs (1.5 mg/mL, 200 μL) via the tail vein, the mouse was immobilized on a lab-built plate and imaged under the three-photon fluorescence microscopic system. The z-stack images were taken at 2-μm step and the scanning speed was 2.2 μs/pixel (512 × 512 pixels per frame).

### *In vivo* three-photon fluorescence microscopic brain imaging of the mouse with intact skull

Animal preparations with intact skull window and optical clearing window followed standard surgery reported previously^39^. The Balb/c mice (male, ∼6 weeks old) with intact skull window were injected with DCBT NPs (1.5 mg/mL, 200 μL) through tail vein. Through-skull imaging was then carried out on the three-photon fluorescence microscopic system. To evaluate the improved effects of imaging depth as well as imaging quality of 3PF microscopic cerebrovascular imaging through the established vis-NIR-II skull optical clearing window, the skull window was treated with S1 and S2 of VNSOCA to get skull optical clearing window. After that, imaging of the same field of view was conducted on the three-photon fluorescence microscopic system. *Thy-1-GFP-M-line* male mice (∼6 weeks old) were treated by the aforementioned operation to obtain the skull optical clearing window. And 3PF neuronal imaging and dual-channel cerebrovascular and neuronal imaging were carried out under the same three-photon fluorescence microscopic system.

### Data analysis

Image J software (Version 1.6.0, National Institutes of Health, USA) was applied for quantitative analysis of each fluorescent image. Origin Pro software (Version 9.0, OriginLab Company, USA), Imaris (Version 9.0, Oxford Instruments) and Adobe Illustrator CC (Version 2018) was applied for graph generation.

## Acknowledgements

This work was supported by National Natural Science Foundation of China (61975172, 82001874, and 21974104) and Natural Science Foundation of Zhejiang Province (LR17F050001).

The authors thank Dandan Song in the Center of Cryo-Electron Microscopy (CCEM), Zhejiang University for her technical assistance on STEM.

## Author contributions

J.Q. and M.H. conceived the idea and designed the experiments. D.L. and D.Z. provided skull optical clearing technique guidance. Z.Z. provided the fluorescence probes. H.Z. and W.X. provided help on 3PM setup system. T.W performed the simulation. L.Z. provided guidance on cranial surgery. Z.H. helped to perform experiment in Fig. 5. M.H., D.L., W.G. Z.H. Z.F. performed the experiments. M.H., D.L., W.G., S.P. and J.Q. analyzed the data and wrote the manuscript. All authors discussed the results and commented on the manuscript.

