## supplementary Figures for "Ultra-deep through-skull mouse brain imaging via the combination of skull optical clearing and three-photon microscopy"

### Supplementary methods

#### Monte Carlo simulation of emission photon penetration through brain tissue and skull

As to the simulation of fluorescence collection, photons emit at random angles from a fixed point. After successfully escaping from the tissue, photon angle will be recorded. A lens is placed on top of immersion media to collect photons those have successfully escaped from the tissue. The radius of the lens is the same as the radius of the focused beam, and the focal length of the lens is calculated to ensure that the distance from the virtual object point in the tissue to the lens is twice of the focal length. A detection plane with 20 mm by 20 mm area is twice of the focal length parallel above the lens. Remaining energy of the photons reaching the detection surface will be recorded. The absorption coefficients of brain tissue and skull were set as 2 $cm^{-1}$ and 0.0046 $cm^{-1}$ in the red-emission channel (referring to the absorption coefficients at 650 nm) respectively. The scattering coefficients of brain tissue and original skull were set as 15.858 $cm^{-1}$ and 18.978 $cm^{-1}$ in the red-emission channel (referring to the scattering coefficient at 650 nm) respectively. Other simulation parameters were consistent with those in Fig. 1.

#### Monte Carlo simulation of NIR photon penetration with USOCA and VNSOCA

There are 3 layers in the Monte Carlo simulation, representing the immersion media of the objective lens, the mouse skull and mouse brain tissue, respectively. The simulation process of focusing light in the tissue is similar to the simulation in Fig. 1. The skull optical clearing induced by USOCA and VNSOCA reduces the skull scattering coefficient, which is set from 100% to 10%. The absorption coefficient of immersion media refers to absorption spectra in Fig. 1i. Other simulation parameters were consistent with those in Fig. 1.

#### *Ex vivo* three-photon fluorescence microscopic imaging

*Ex vivo* test was carried out as following: Firstly, the skull of a mature female mouse was removed and cued on the plate. Then the glass capillary tube with DCBT NPs was placed beneath the skull. With $D_2O$ covered, the *ex vivo* through skull imaging was carried out under three-photon fluorescence microscopic system. After that, the skull was treated with solution 1 (S1) for 10-15 min. Then, solution 2 (S2) was used to replace S1 when the skull clearing was established. After all the procedures, the capillary tube beneath the optical transparent skull was imaged again under three-photon fluorescence microscopic system. The clearing effect was demonstrated by contrasting three-photon fluorescence images of DCBT NPs before and after skull clearing treatment. The distance between the skull and capillary tube could be adjusted to simulate different imaging depths *in vivo*.

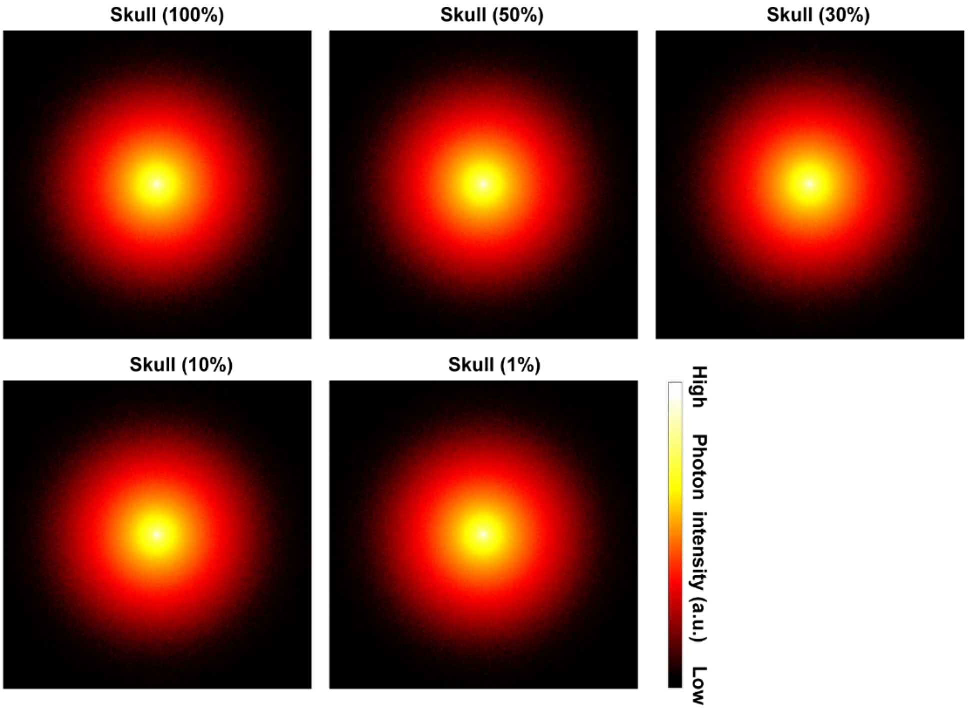


**Supplementary Figure 1.** X-Y plane photon distribution stimulation of red-channel (650 nm) fluorescence signal at the skull surface, which was three-photon excited and then penetrated through brain tissue and skull with different remaining skull, scattering ratio (100% to 1%).

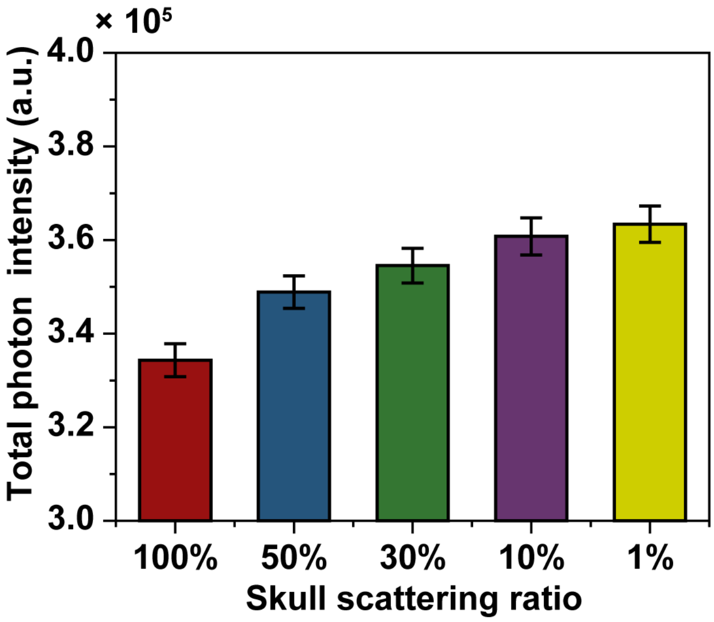


**Supplementary Figure 2.** Total photon intensity of 3PF at skull surface in supplementary Figure 1 with different remaining skull scattering ratios (100% to 1%).

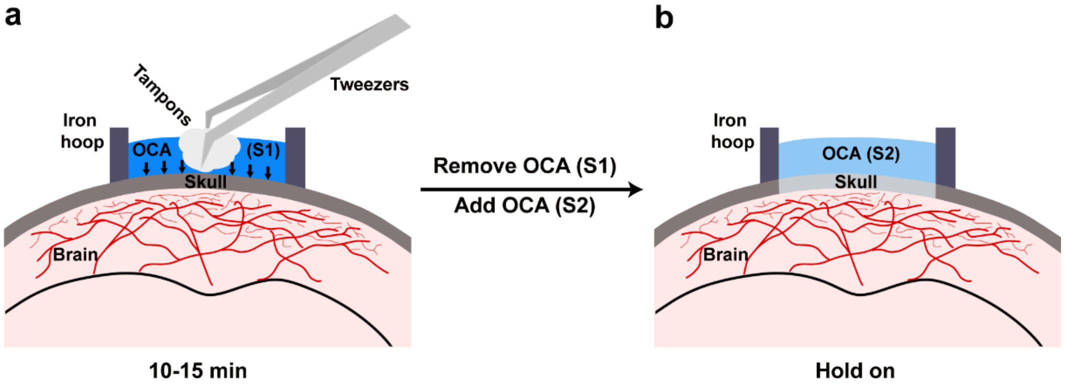


**Supplementary Figure 3. Schematic of skull optical clearing procedure. a** The skull was dealed with solution 1 of optical clearing agents for 10-15 min. The tampons were applied to press the solution 1 to seep down into the skull. **b** The solution 2 of optical clearing agents was added to replace solution 1 for the subsequent 3PF imaging.

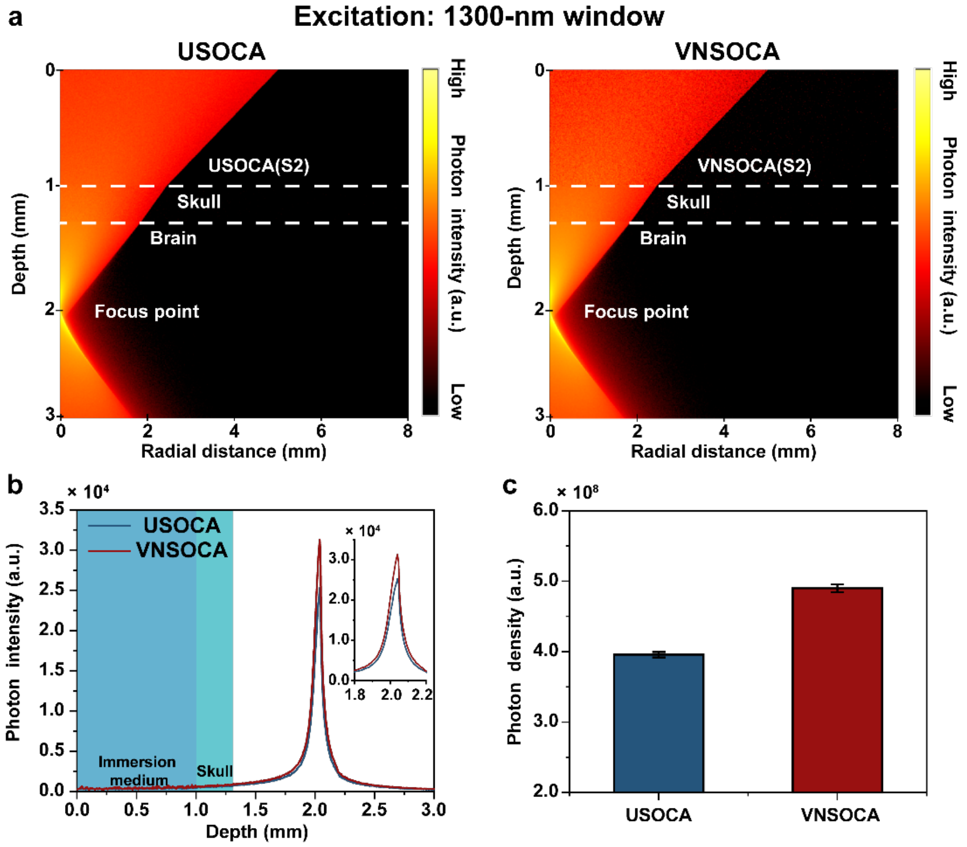


**Supplementary Figure 4. Photon distribution stimulation of excitation light in the 1300-nm window. a** R-Z plane photon distribution stimulation of excitation light in the 1300-nm window penetrating through immersion medium, skull and brain tissue with two different skull optical clearing agents. **b** Photon intensity profiles along the depth (Radial distance = 0 mm) in (a). **c** Photon density at the focused point with different skull clearing agents in (a).

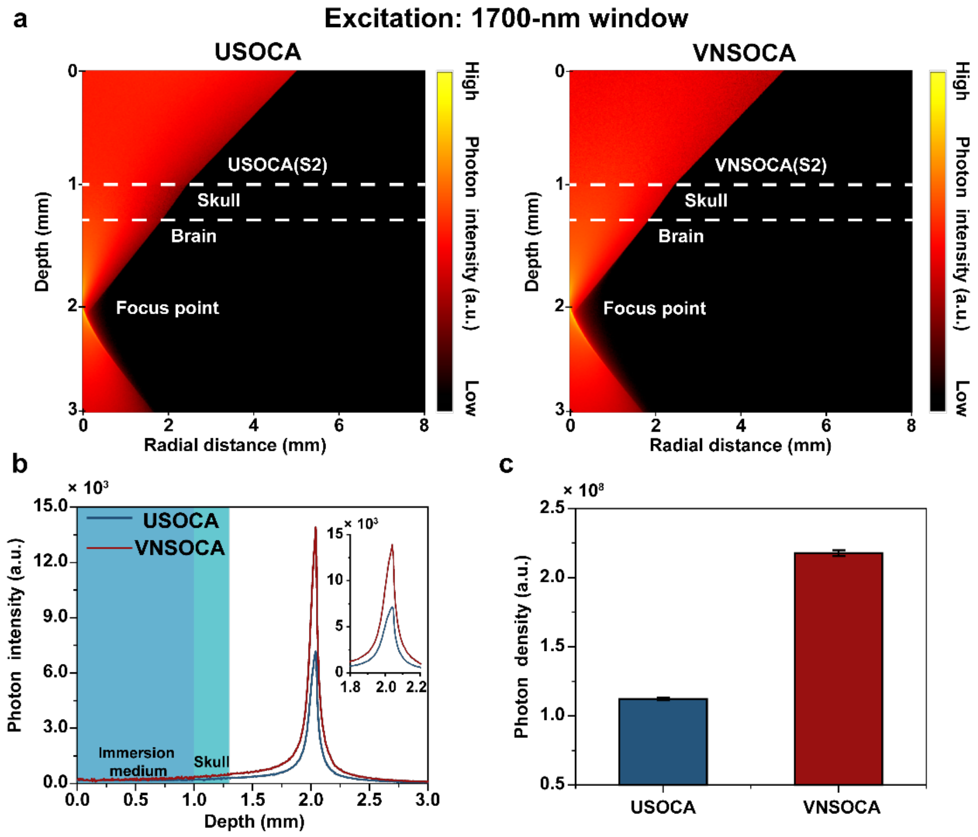


**Supplementary Figure 5. Photon distribution stimulation of excitation light in the 1700-nm window. a** R-Z plane photon distribution stimulation of excitation light in the 1700-nm window penetrating through immersion medium, skull and brain tissue with two different skull optical clearing agents. **b** Photon intensity profiles along the depth (Radial distance = 0 mm) in (a). **c** Photon density at the focused point with different skull clearing agents in (a).

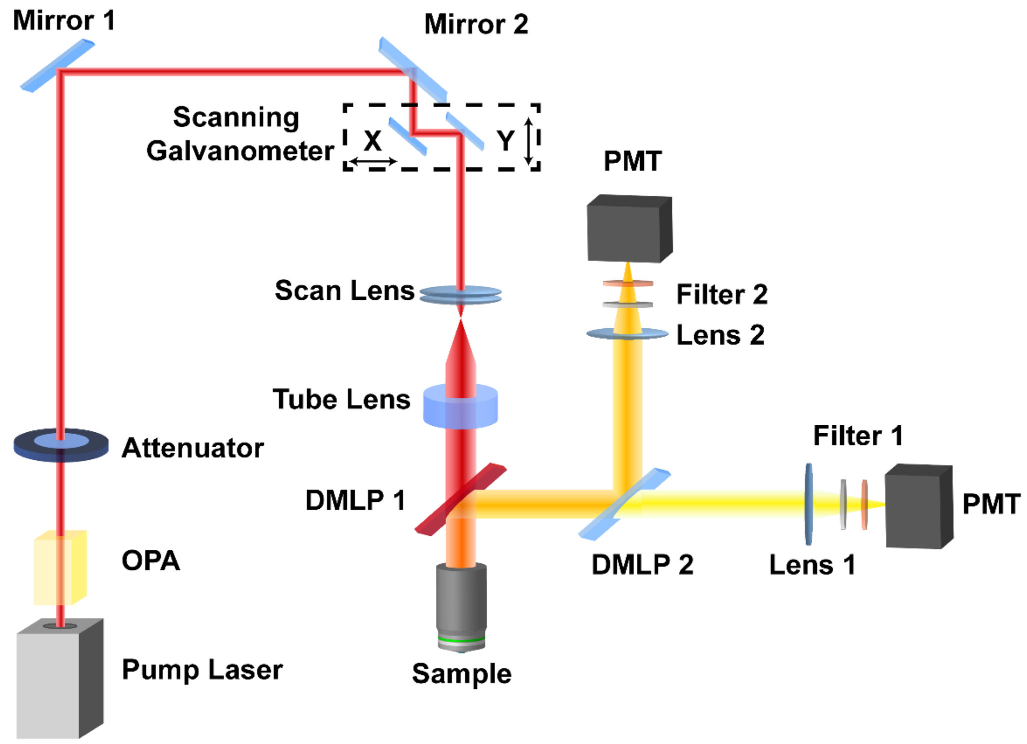


**Supplementary Figure 6.** Schematic illustration of the three-photon microscopic system.

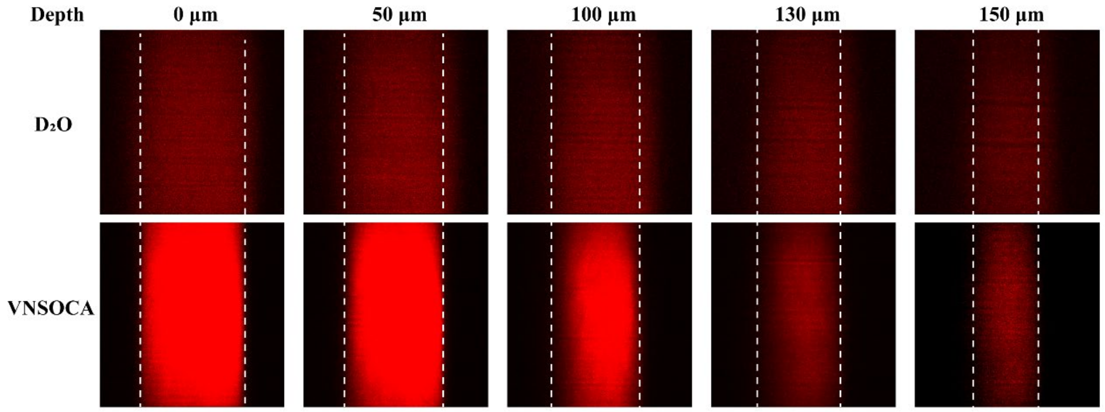

**Supplementary Figure 7.** 3PF imaging of the DCBT NPs-filled capillary covered by a mouse the skull before and after skull optical clearing at different focus plane. Excitation condition: 1300 nm fs laser.

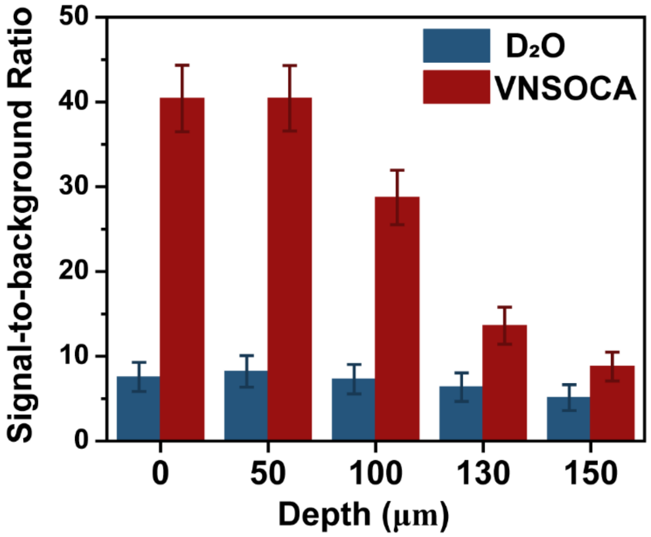


**Supplementary Figure 8.** The signal-to-background ratio (SBR) analysis of the 3PF imaging results in supplementary Figure 7.

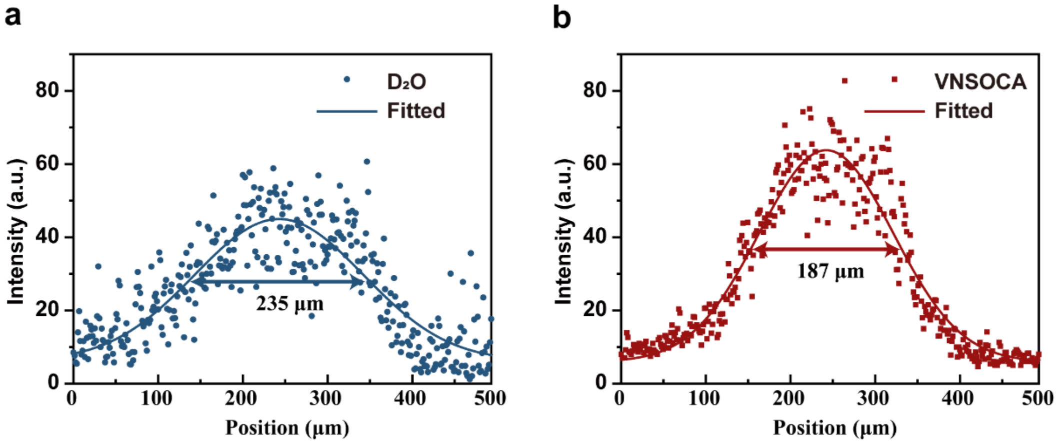


**Supplementary Figure 9.** Full Width at Half Maximum (FWHM) analysis of 3PF imaging of DCBT NPs before (a) and after (b) skull optical clearing at 150 μm depth, in supplementary Figure 7.

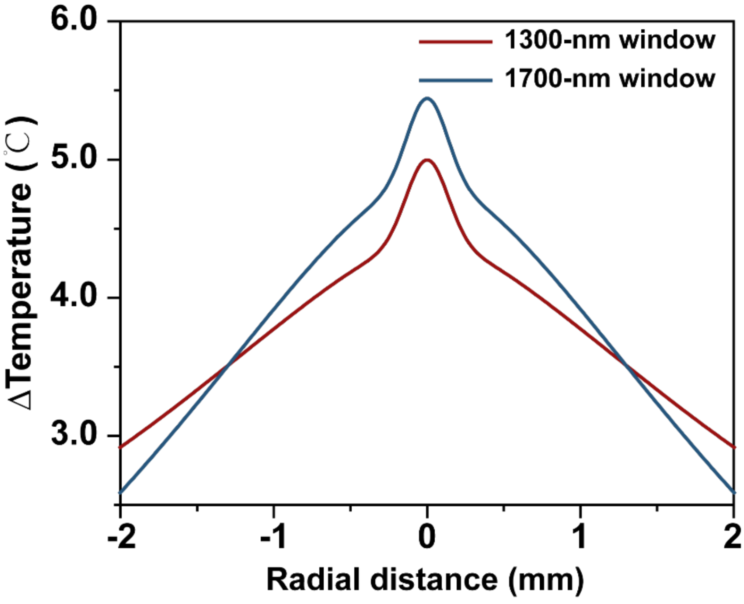


**Supplementary Figure 10.** Temperature rising distribution line versus radial distance at the position of 1 mm focal depth in Fig. 2a.

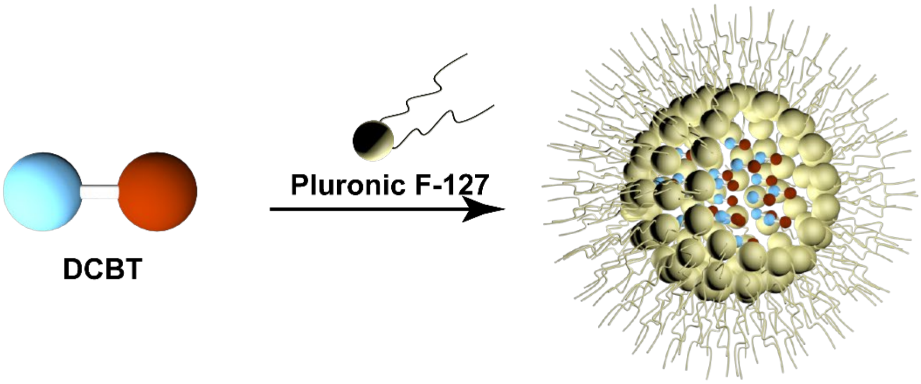


**Supplementary Figure 11.** Schematic illustration of the fabrication of DCBT NPs.

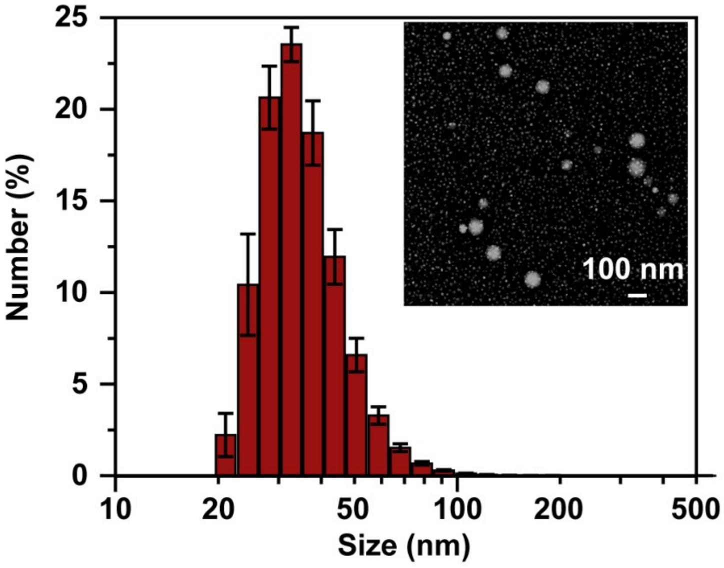

**Supplementary Figure 12.** DLS results of DCBT NPs and a representative TEM image of DCBT NPs.

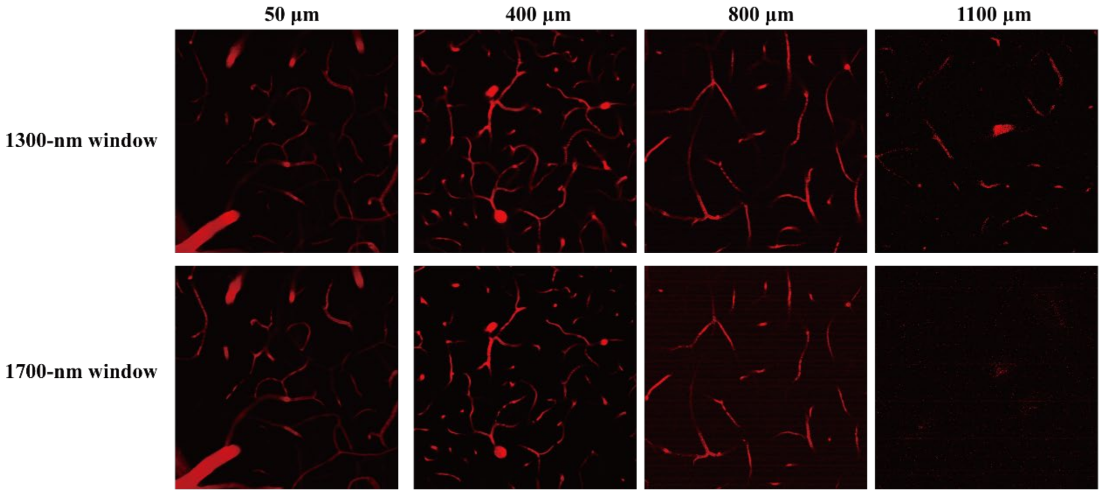


**Supplementary Figure 13.** 3PF imaging of the mouse brain vasculature at various depths excited using the 1300-nm and 1700-nm window in Fig. 3.

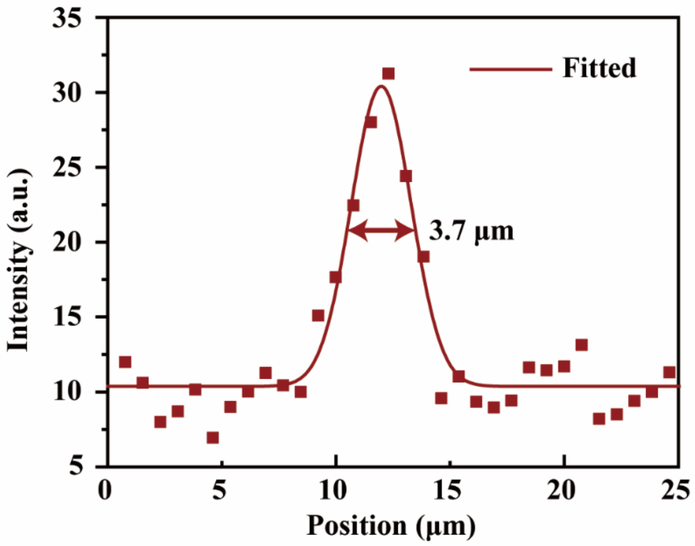


**Supplementary Figure 14.** Intensity and FWHM analysis of a blood vessel at the depth of 1500 μm in Fig. 3j.

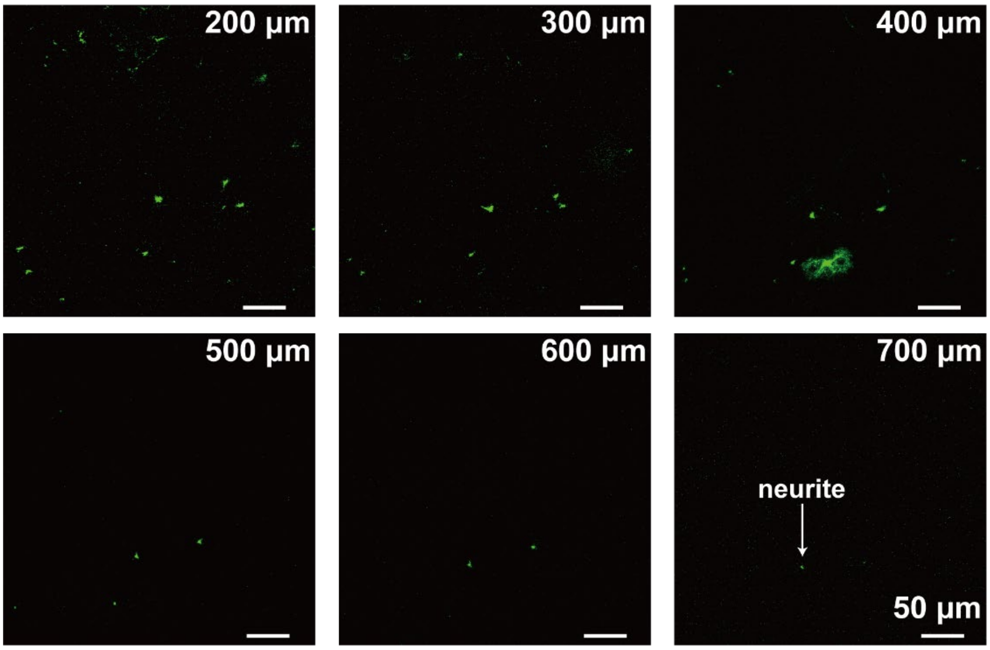


**Supplementary Figure 15.** 3PF deep imaging of the mouse brain neurons after skull clearing in Fig. 5.
